# ARDUINO-BASED ORGAN PERFUSION SYSTEM

**DOI:** 10.1101/2025.01.13.632705

**Authors:** Stefano Di Domenico

**Author notes:** **Author contributions:** SDD conceived the project, built the devices, wrote the programs, conducted the calibration and the statistical tests, and wrote the article.

## Abstract

Laboratory equipment could be expensive and their cost can be unsustainable for scientists with limited financial resources. In order to overcome these impediments and to improve our experimental studies on liver resection in rats, a multipurpose organ perfusion system was projected and realized using low-cost components.

The device is based on Arduino boards, an easy-to-use and open-source microcontroller, that receives digital and analog inputs, can produce digital outputs and it is programmable in C++. Our perfusion system is composed of: 1) the main control unit that receives pressure data from two independent pressure transducers commonly used in the OR, it receives input from two potentiometers, seven switches, and a push-button; it can display data with an LCD and it can control the speed and the direction of two independent peristaltic-pump, 2) a motor unit composed by peristaltic pumps controlled by two stepper motors, 3) a thermostatic unit that received data from temperature probe and can control up to three heating units.

The pulsating flow generated by the peristaltic pump can be flattened by a dampener unit to simulate the portal flow, and the interposition of a hollow-fiber micro-oxygenator can increase the pO2 of the perfusion fluid in the arterial line.

This cheap system is designed to measure the compliance of regenerating rat liver at increasing and decreasing portal and arterial flow during normothermic ex-vivo perfusion. However the system can be easily rearranged to perform different kinds of organ perfusion: subnormothermic o hypothermic ex-vivo preservation tests can be done by adding a Peltier-based cooling module. Indeed the main control unit can be programmed to control the flow rate based on pressure data to realize peritoneal perfusion or isolated limb perfusion.

## BACKGROUND

Isolated perfusion has represented for decades the best model for studying the physiology and pathophysiology of many organs [1,2].

Technologies developed in these field have subsequently allowed the development of heart-lung machine, and chemo-hyperthermic treatments through isolated perfusion of the limbs and the peritoneal cavity [3,4].

More recently, isolated organ perfusion is gaining a fundamental role in organ preservation and recondition for transplantation [5,6].

The renewed clinical interest on isolated organ perfusion accounts for the exponential development of basic and translational research lines aimed at studying the complex relationships between organ functionality, hemodynamics, type of perfusion and use drugs during ex-vivo preservation.

However, the perfusion systems on the market require huge funds and have an intrinsic limit of versatility, limiting the opportunities for researchers with scarce financial resources to undertake new studies and test new hypotheses.

To overcome these limitations, I designed and built an open-source, scalable and low-cost organ perfusion system for research purposes, based on Arduino boards, cheap materials, and reverse-engineering commercial sensors.

## MATERIALS AND METHODS

The basic perfusion system comprises a power supply, the main control unit, the peristaltic pump unit, the pressure transducers, the dampener, the oxygenator and the thermostatic unit (Figure 1).

**Figure 1.**
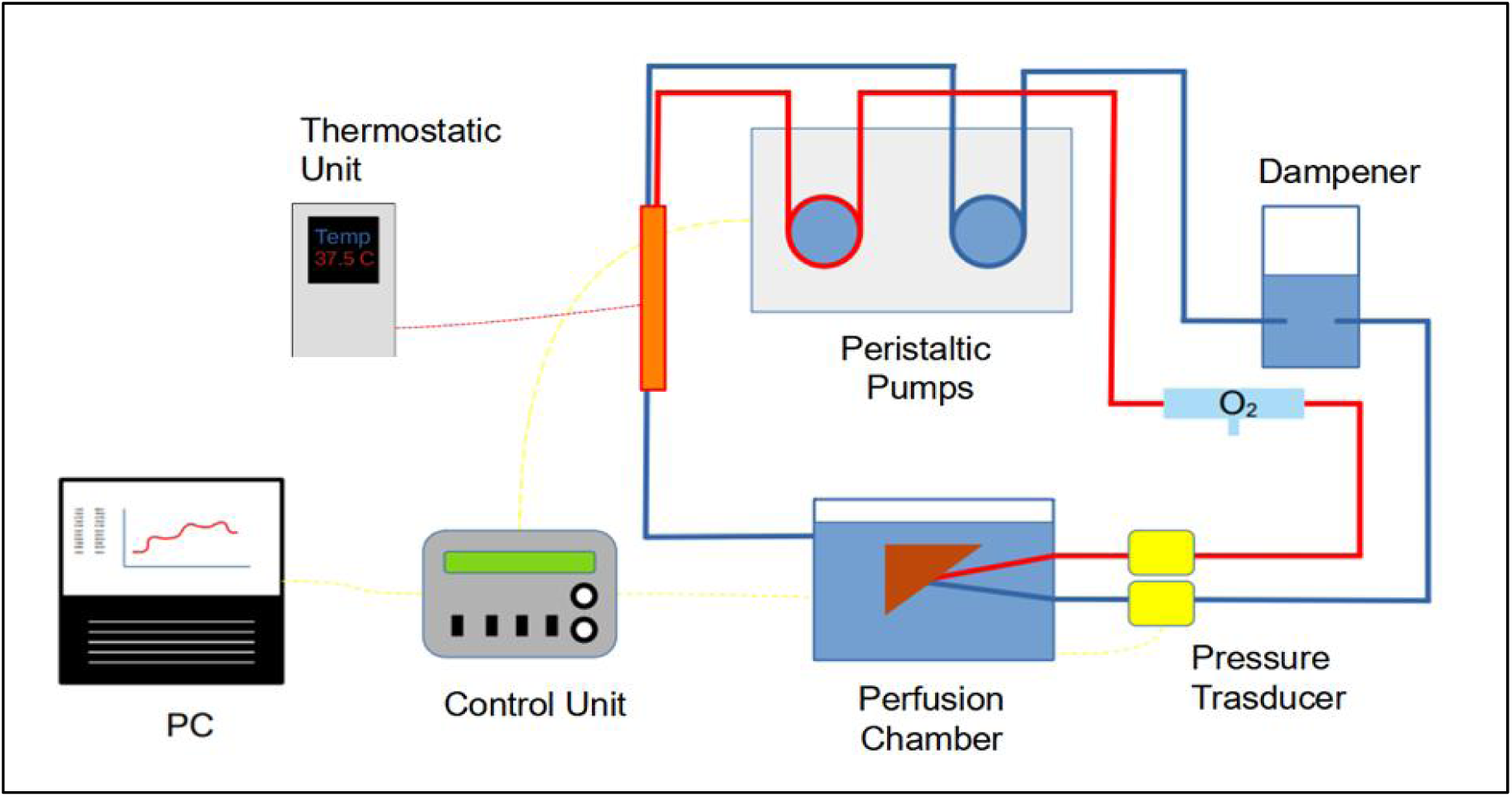
Components of the organ perfusion system.

### The power supply

A commercial 220V AC to 12V DC 10A switching power supply is connected to a digital voltmeter amperemeter and encased in a junction box (Figure 2).

**Figure 2.**
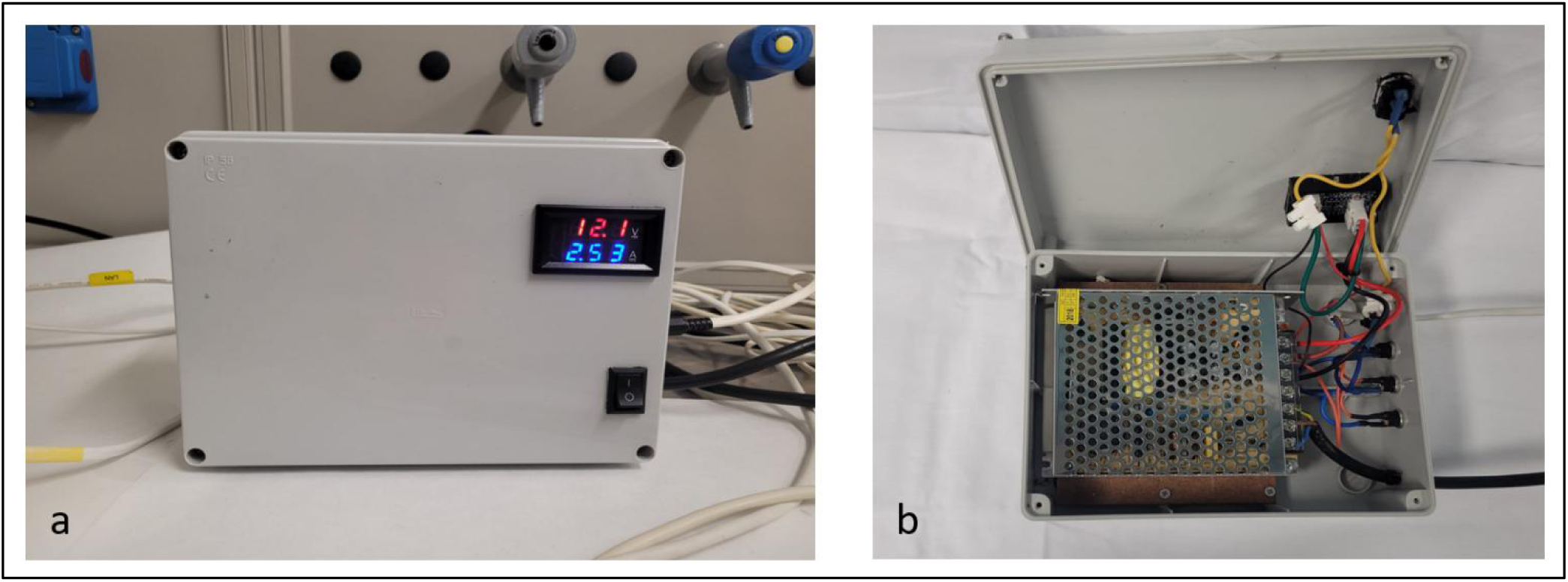
The power supply with the main switch and voltmeter-amperometer (a); the internal view with the DC 12V power switching and the three output jacks (b).

It supplies the main control unit with a 5.5 x 2.5 mm plug and it can also deliver electricity by additional three jacks.

### The main control unit

Arduino board is a family of open-source, low-cost, integrated circuits that contain an Atmel AVR processor, memory, and analog and digital input and output pins. These boards can be programmed in C++ within the Arduino Integrated Development Environmental (IDE) to acquire inputs from a variety of switches and sensors and to control lights, motors, and other physical outputs [7].

In short, the main unit is built around an Arduino Mega board, which gathers pressure data from two amplifiers connected to commercial pressure transducers. It also controls the peristaltic pump unit using signals from two Arduino Nano boards. The unit can be controlled using analog inputs from a potentiometer, digital inputs from five toggle switches, and one push button. Additionally, it can display data using an LCD (Figure 3).

**Figure 3.**
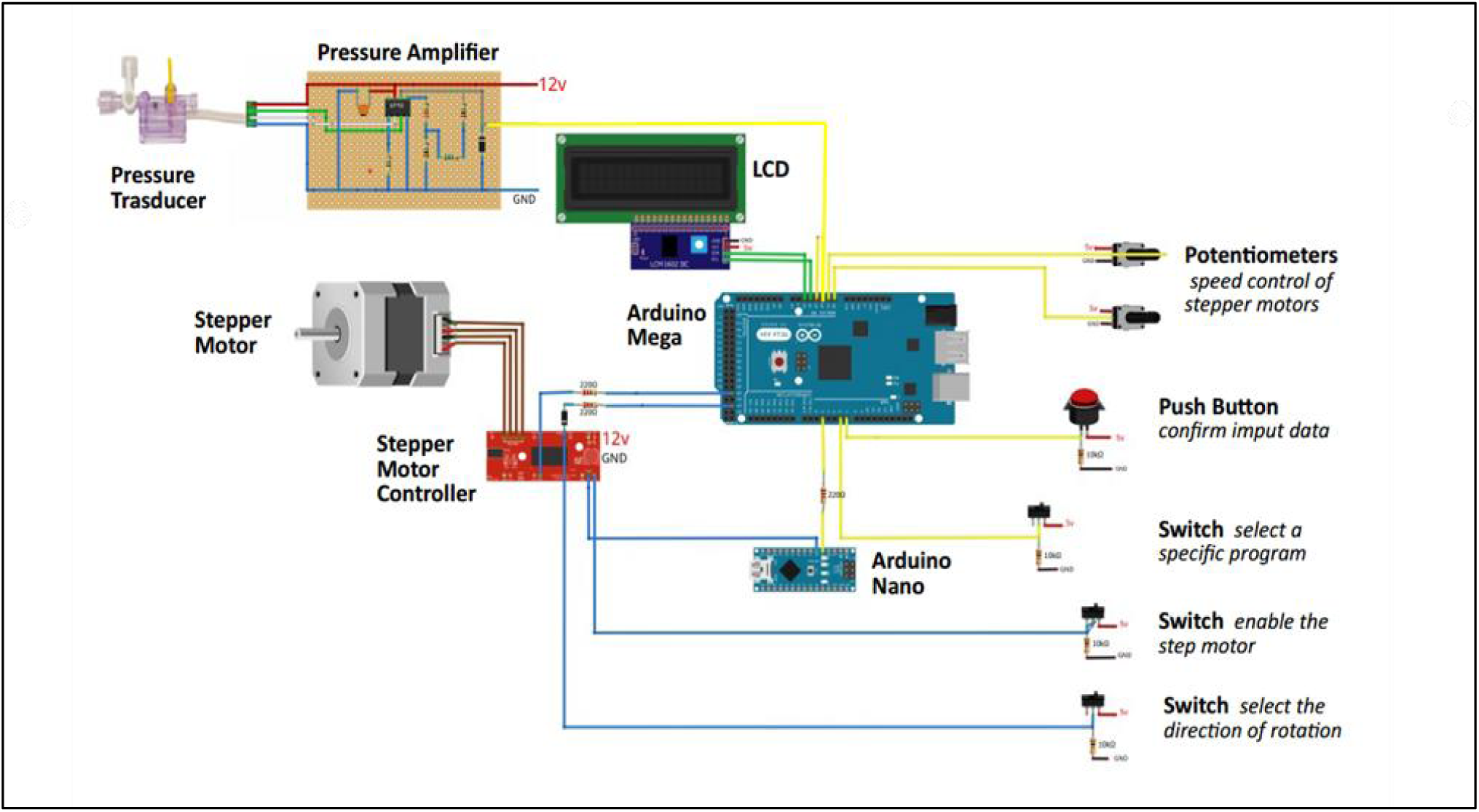
Wiring diagram of the control unit: the actual module is based on two stepper motors and two pressure transducers, but in the figure, only one of each is shown.

More in details, in this project an Arduino Mega board is powered by DC 12V from the DC power supply, and it receives pressure analog signals from two amplifiers varying from 0 to 4.8 volts that are converted from 0 to 1023 digital values with a resolution of 4,6875 mV.

It receives two distinct analog inputs varying from 0 to 5 volts from two 10 Kohm potentiometers (4,88 mV resolution) that are used to assign specific values to variables in the program, such as peristaltic pump velocity, organ weight, or perfusion time interval.

It received a digital input from a push bottom and seven toggle switches, connected through 10 Kohm pull-down resistors, that are used to choose between discrete options of the program, to directly enable the stepper motor and to select the sense of rotation of the peristaltic pump.

It produced two distinct pulse-width modulation (PWM) signals with a frequency of 980Hz and duty cycle varying from 0% to 100% that are sent converted to a It produced two distinct pulse-width modulation (PWM) signals with a frequency of 980Hz and duty cycle varying from 0% to 100% that are sent to two Arduino-nano boards. The PWM signals received by the Arduino nano is then converted to square wave signals with variable frequency between 0 and 2020 Hz that are sent to the motor unit to control the velocity of the peristaltic pumps.

An LCD display was used to visualize some crucial information such as the states of the program and the value of some variable: it is able to display 16 characters arranged in 2 rows, and it is controlled by the Arduino board with an I2C module using only two digital pins.

The main control unit components were encased in a recycled box from a dismissed glucometer (Figures 4-5).

**Figure 4.**
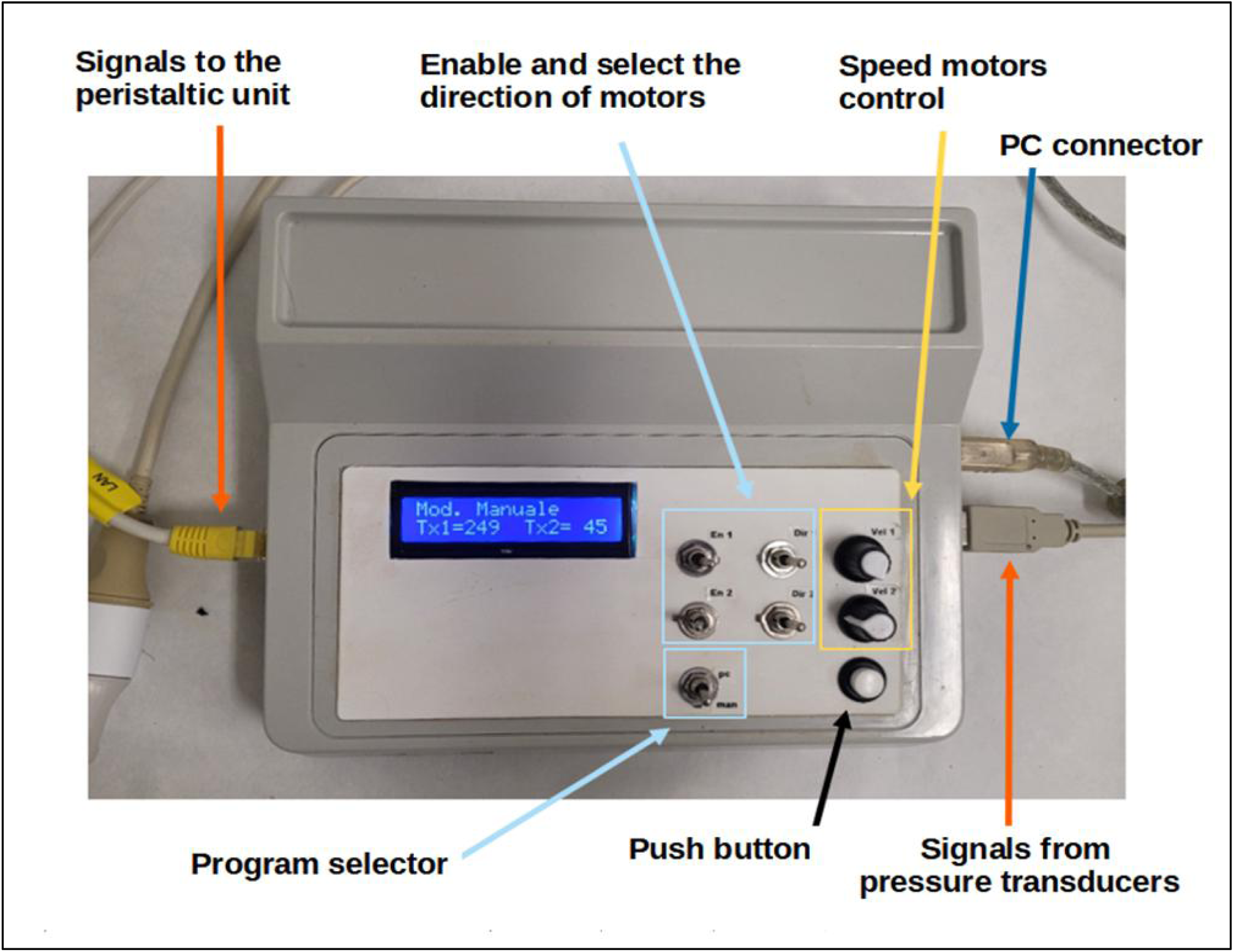
The control unit with two potentiometers, a push button, toggle switches, the LCD and the cables connected.

**Figure 5.**
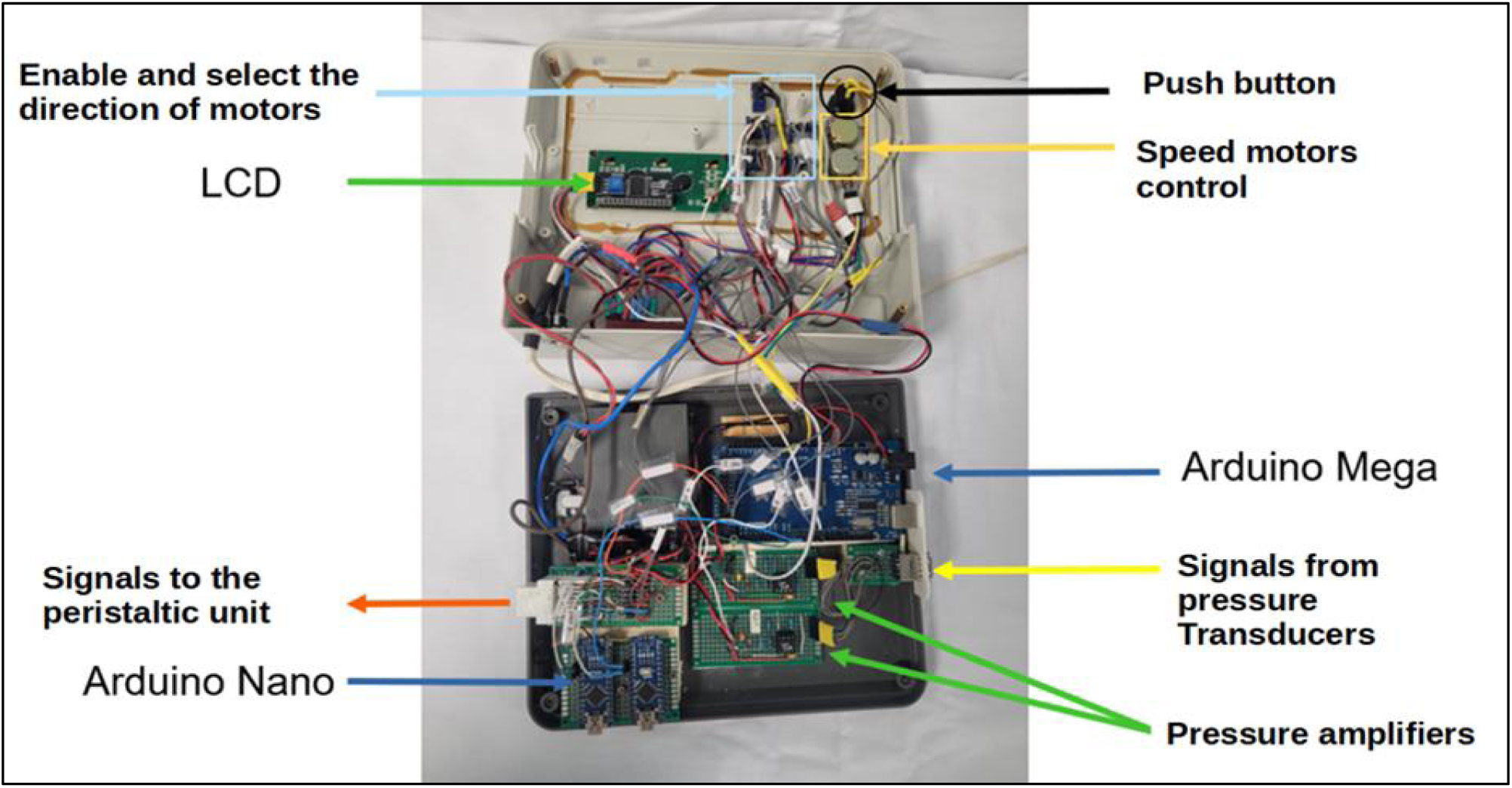
The modular elements inside the control unit are visible.

Digital inputs and outputs can be visualized and recorded in any PC through an USB port using CoolTerm, a freeware serial port terminal application that establishes a connection between computer and external devices in order to exchange and analyze data. It is freely available at http://freeware.the-meiers.org.

### Pressure Transducer

TruWave is a disposable pressure transducer distributed by Edwards Lifesciences® that communicates blood pressure data from a pressure catheter to patient monitoring systems [8]. It is widely used in operative rooms and intensive care units. Despite its main specifications are available on the producer website, the construction details are patent-protected, and some assumptions have been made and tested in order to use it for our purposes as previously described [9].

The TruWave sensor output μV (10^-6 volts) must be amplified to mV (10^-3 volts) range in order to be clearly detectable by the microcontroller-board, since its resolution is around 4,9 mV (5V / 1024).

Two simple amplifiers were built for this purpose, using the OP90 integrated circuit obtained from a discarded glucometer. Following the instructions in its data sheet [8], two single-supply instrumentation amplifiers were assembled. Different sets of resistors were tested to achieve the desired pressure range amplification : 0-40 mmHg and 0-150 mmHg (Figure 6).

**Figure 6.**
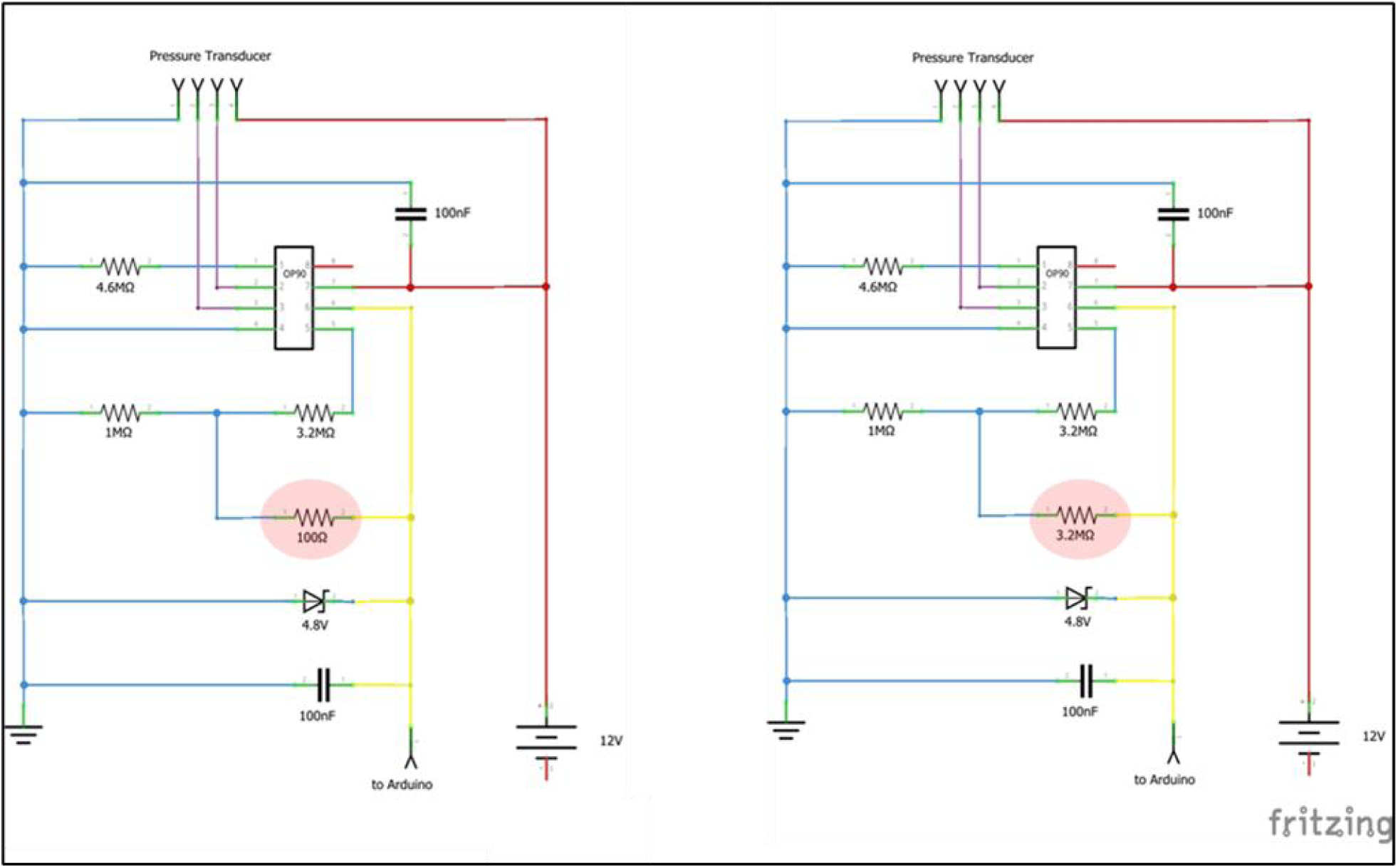
Schematic of the two amplifiers based on OP90 for arterial pressure range [0-145 mmHg] on the left, and for portal venous pressure range [0-45 mmHg] on the right: the highlighted resistances set the gain.

### The peristaltic unit

The peristaltic unit is a system made of two peristaltic pumps driven by two 42-step stepper motors, powered at 12v DC and controlled by two Easy-Driver boards (Figure 7).

**Figure 7.**
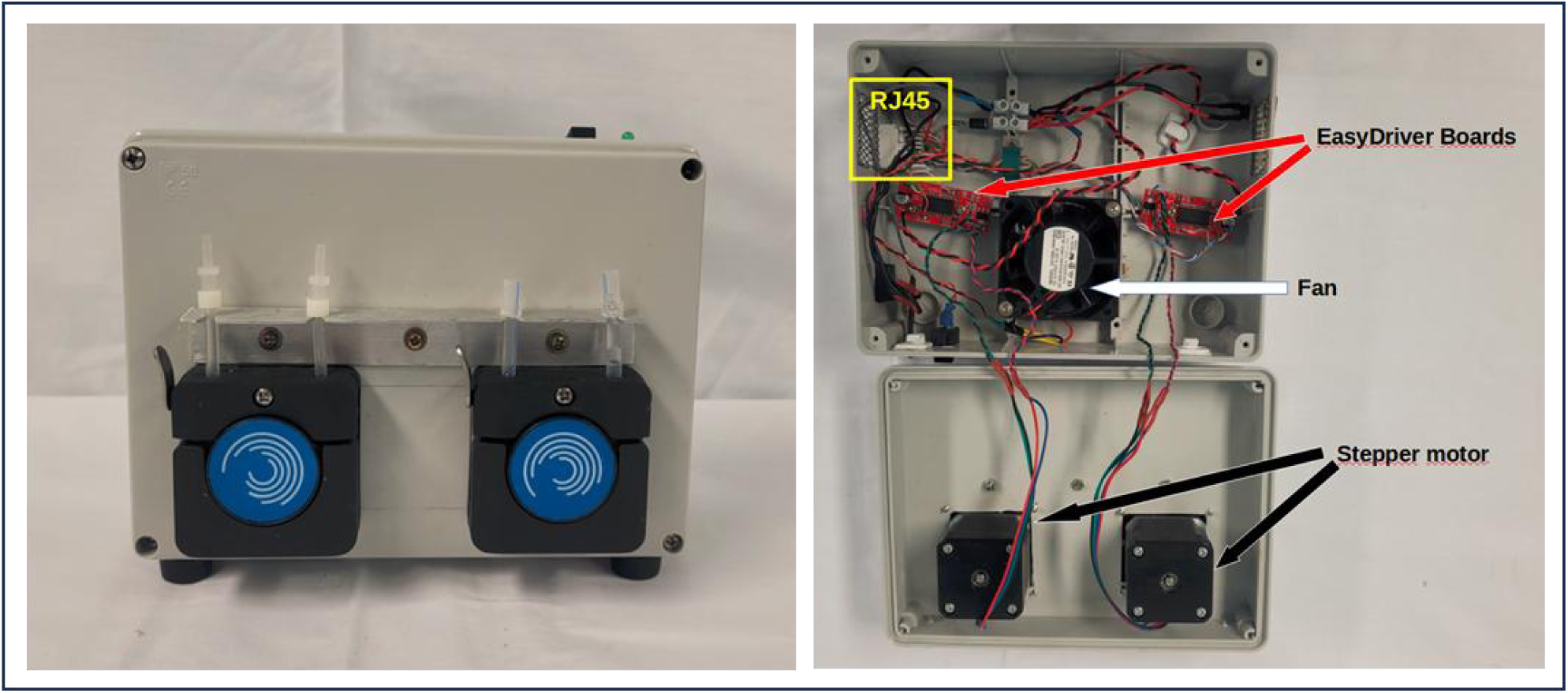
On the left is the external appearance of the two peristaltic pumps with different tube sizes; on the right is the inner aspect with the fan, the EasyDriver boards and the connector.

Each Easy-Driver boards receive from the control unit a 0-5v logic signal for motor enable, a 0-5v logic signal for the direction of rotation, and a 0-5v square wave signal with duty-cycle 50% and frequency 10-5000Hz for rotational speed control.

The perfusion unit features a 12v fan for cooling the motors and boards and it is encased in a junction box .

The 12v power supply and the digital signals from the control unit are transmitted through an Ethernet cable with RJ45 connectors.

Tubes within the pump compartment has been sized to achieve a flow rate of 0-50 ml/min for portal perfusion and 0-15 ml/min for arterial perfusion.

### The dampener

Peristaltic pumps create a pulsing flow that can simulate arterial flow, while a non-pulsating stream, like venous/portal flow, can be obtained using a dampener. A 100ml glass airtight container filled with 50ml of perfusate is used to flatten the peristaltic flow (Figure 8).

**Figure 8.**
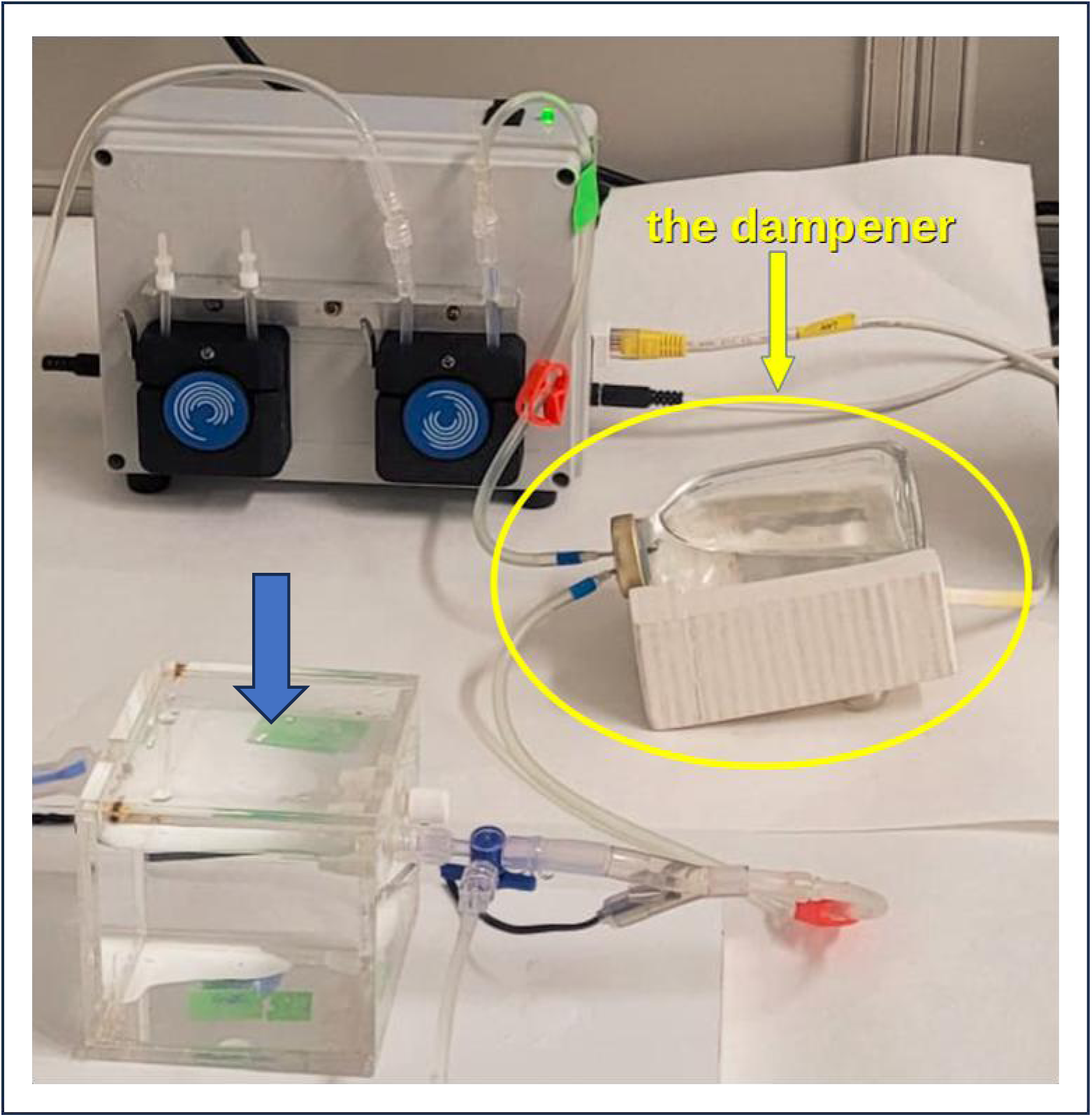
The dampener is connected between the peristaltic pump and the perfusion chamber (blue arrow).

### The basic temperature control unit and the perfusion chamber

The basic temperature control unit consists of a DC 12V programmable digital thermostat with a temperature sensor and power output jacks to supply resistance loads. A single heating wire (R = 30 ohms) from a hairdryer is used to create a heating element. This wire is wrapped around a glass tube and insulated with heat shrinkable tubing. The temperature control unit is enclosed in a recycled laboratory instrument case (Figure 9).

**Figure 9.**
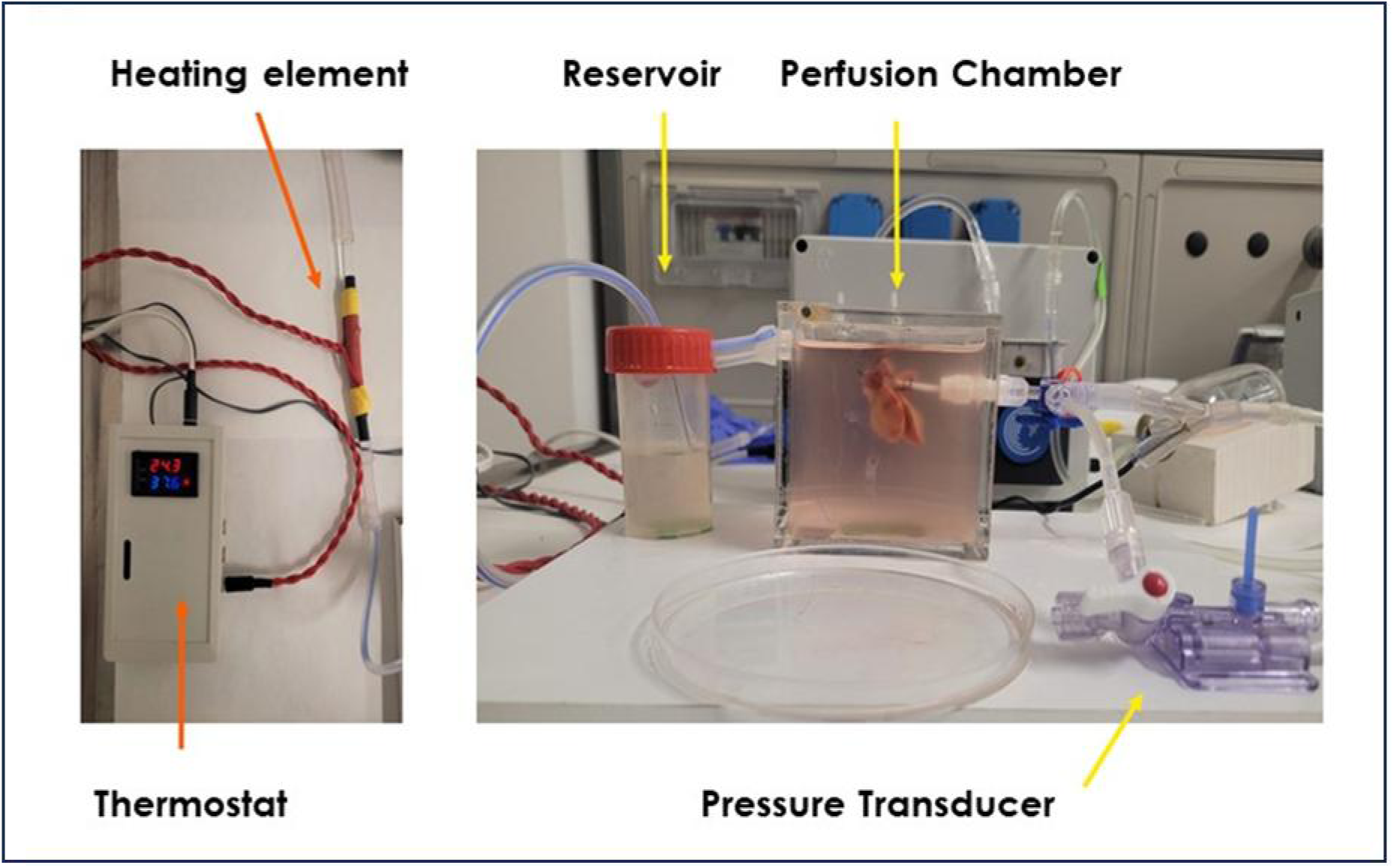
The left side displays the heating element and the thermostat. On the right, a rat liver is floating under normothermic condition in a perfusion chamber connected to the reservoir and pressure transducer.

In a similar way, a second type of heating element is build to hold and warm a 100ml reservoir (Figure 10).

**Figure 10.**
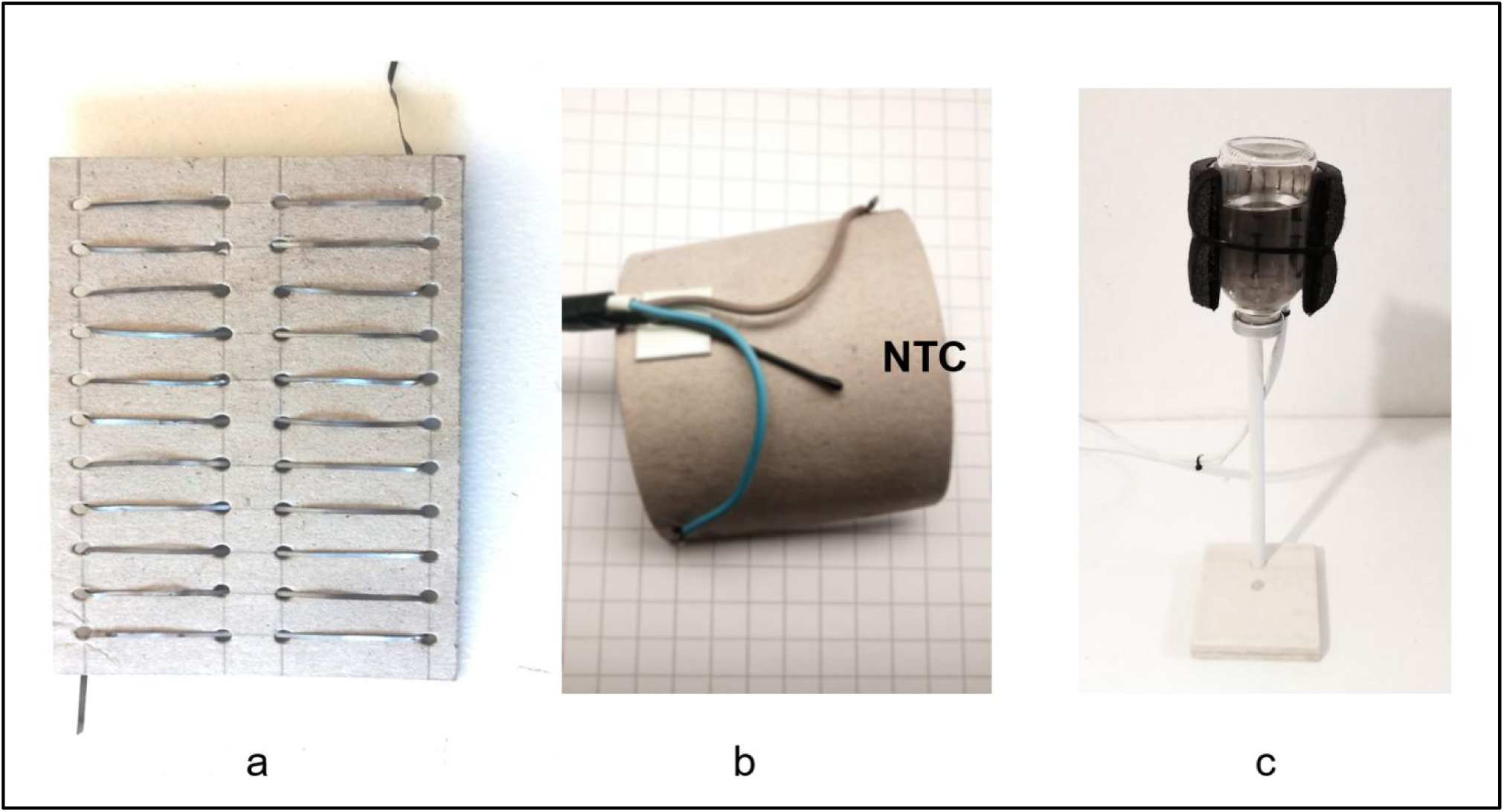
The heater element is build using a resistive wire (a). A thermocouple (NTC) probe is enclosed to monitor and avoid dangerous temperature inside the heater (b). A 100ml airtight bottle is used as perfusate reservoir (c).

### The perfusion chamber

A plexiglass box is used as a perfusion chamber. Two drilled holes serve as inflow using standard luer-lock connectors, while a third hole is used as an outflow connected to a reservoir (Figure 9).

### The oxygenator

The mini-oxygenator is constructed using hollow fibers obtained from an expired oxygenator system. The housing is made up of two drilled Plexiglas blocks sealed with two-component adhesive and containing the hollow fibers. The oxygen travels inside the fibers, while the perfusion fluid flows around them (Figure 11).

**Figure 11.**
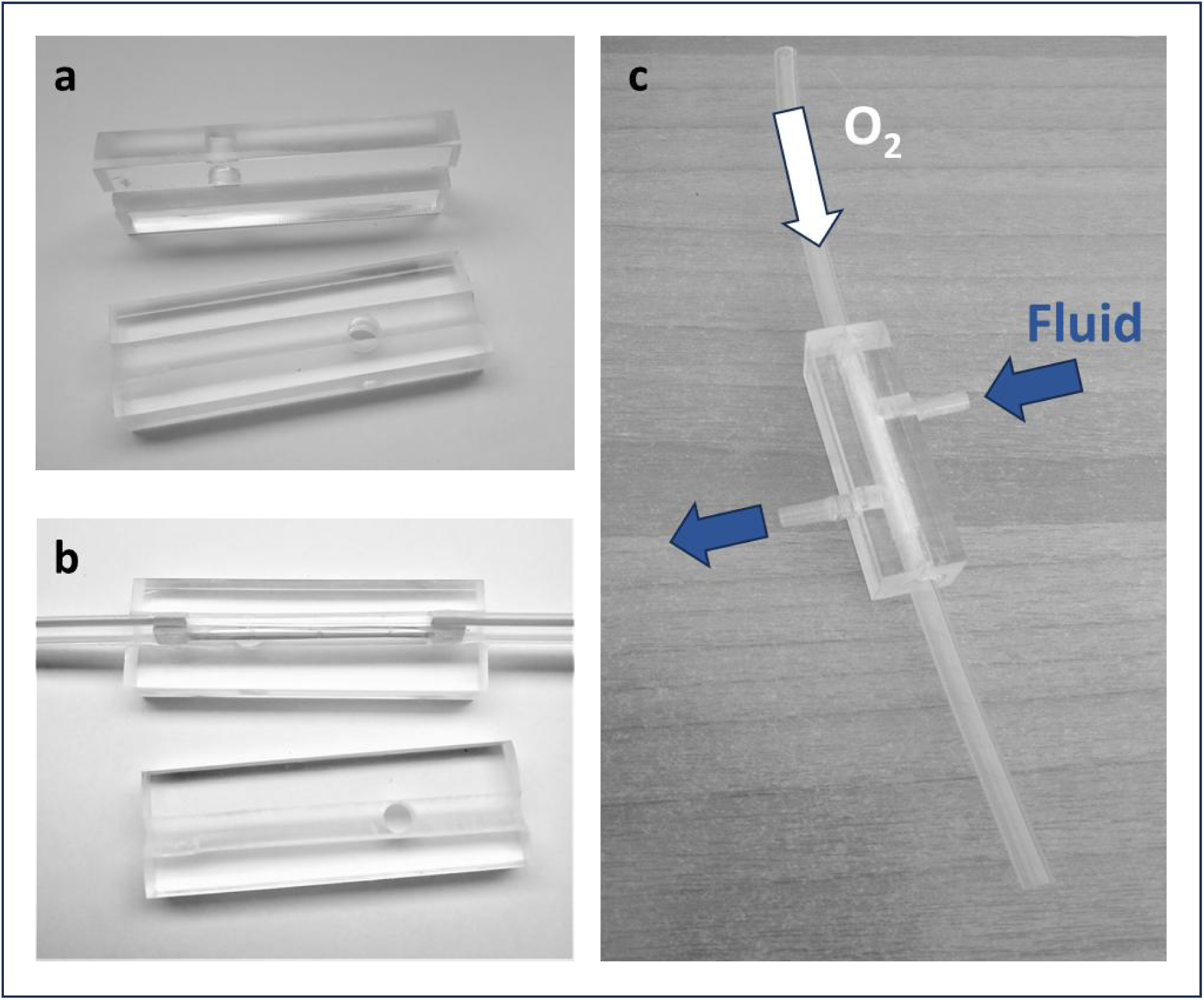
Two drilled plexiglas blocks (a) housed the hollow fibers that are sealed and connected to silicon tubes (b). The direction of fluid and oxygen are displayed in the final product (c).

### Peristaltic pump calibration

The calibration process involved using a precise scale (0.01g) to measure the weight of demineralized water pumped for each Pulse Width Modulation (PWM) value over a fixed period of 5 minutes. Two different sizes of silicon tubes were used for this measurement. The water density was assumed to be 1 g/ml. Specific programs were coded to control the PWM and time.

### Pressure transducer calibration

For this project, we conducted calibration using a water column as the standard, as described earlier [8]. We established the correlation between pressure and digital input values by taking 250 measurements for every 5 cm of water, ranging from 0 to 40 cm. We then converted the values, assuming that 1 cm of water is equal to 0.736 mmHg.

### Statistics

Correlation test, and regression analysis were performed using IBM SPSS Statistic v.22.

## RESULTS

### Pressure transducers

In this project, the measurement rate for each sensor is 25Hz. Both arterial and venous pressure transducers show a linear correlation, with statistical significance at Pearson test with a Spearman Rho equal to 1 (Figure 12, 13).

**Figure 12.**
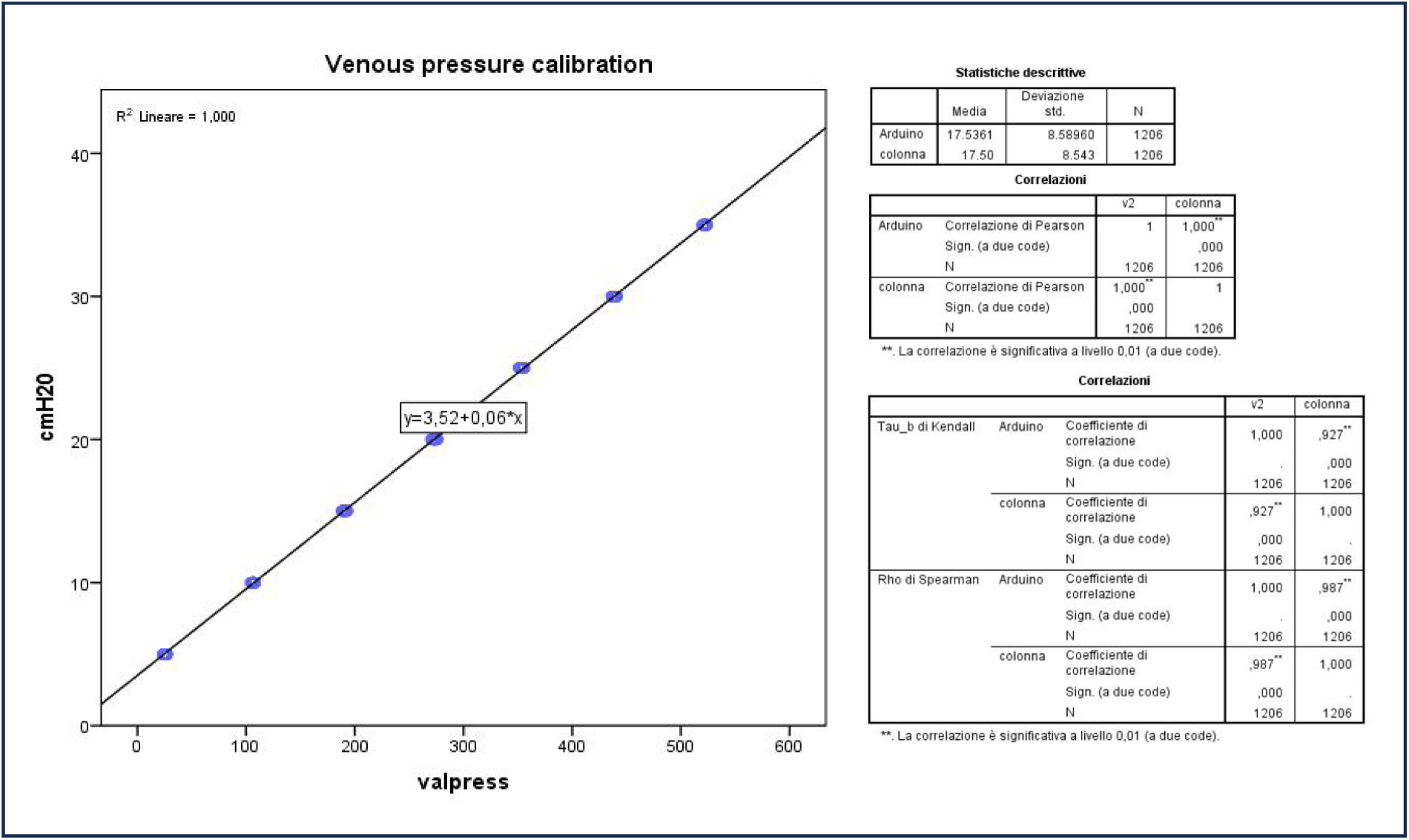
Correlation between the venous pressure values (cmH2O) and the value registered by Arduino board (valpress).

**Figure 13.**
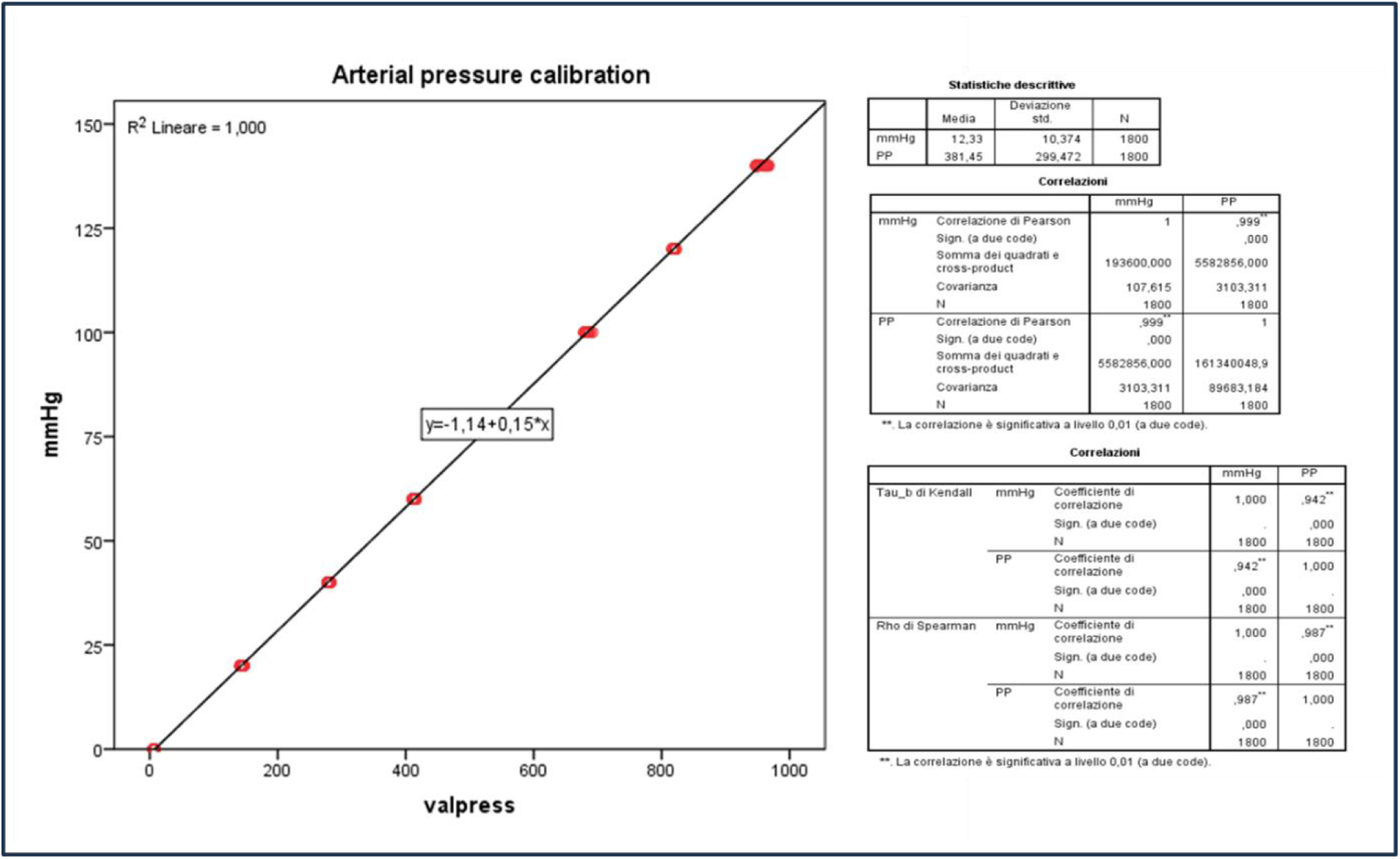
Correlation between the arterial pressure values (mmHg) and the value registered by Arduino board (valpress).

### The Dampener

The dampener effectively reduced the pulsatile flow from the peristaltic pump. At faster rotational speeds, the dampener is capable of flattening pressure variations tenfold (Figure 14).

**Figure 14.**
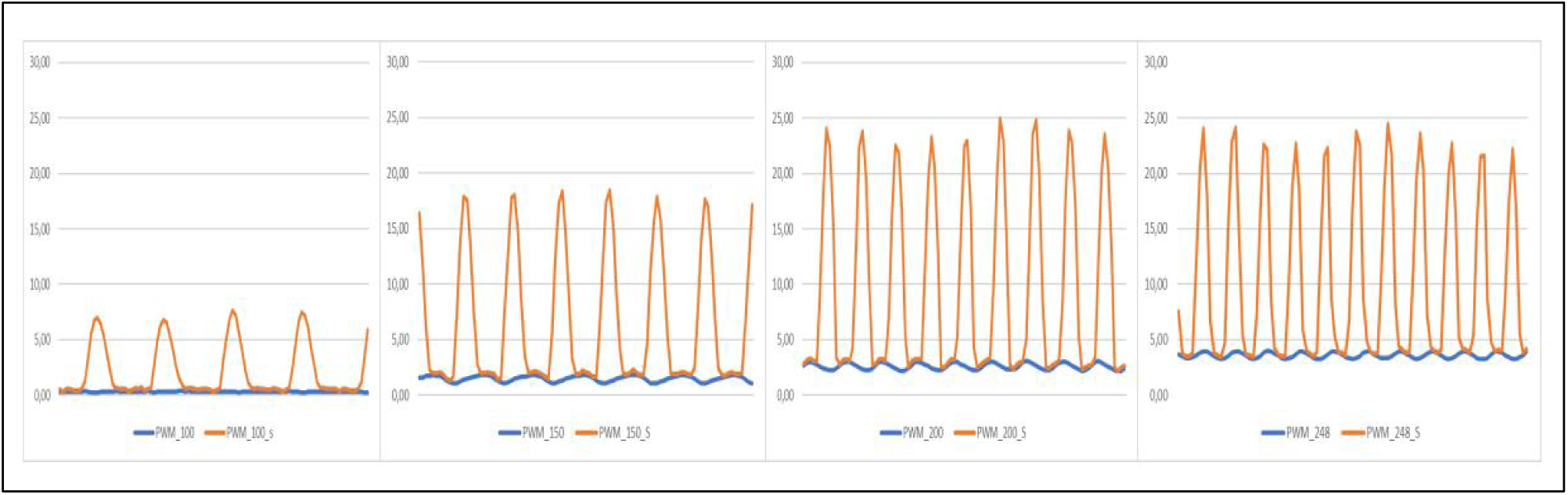
The peristaltic pump produces peristaltic flows (orange lines) that are attenuated by the dampener (blue lines) at each rotational speed controlled by the PWM signals.

### The peristaltic unit

The tubes within the pump compartment are sized to achieve a flow rate of 0-50 ml/min for portal perfusion and 0-15 ml/min for arterial perfusion. Both arterial and venous flow rate show a linear correlation, with statistical significance at Pearson test with a Spearman Rho equal to 1 (Figure 15).

**Figure 15.**
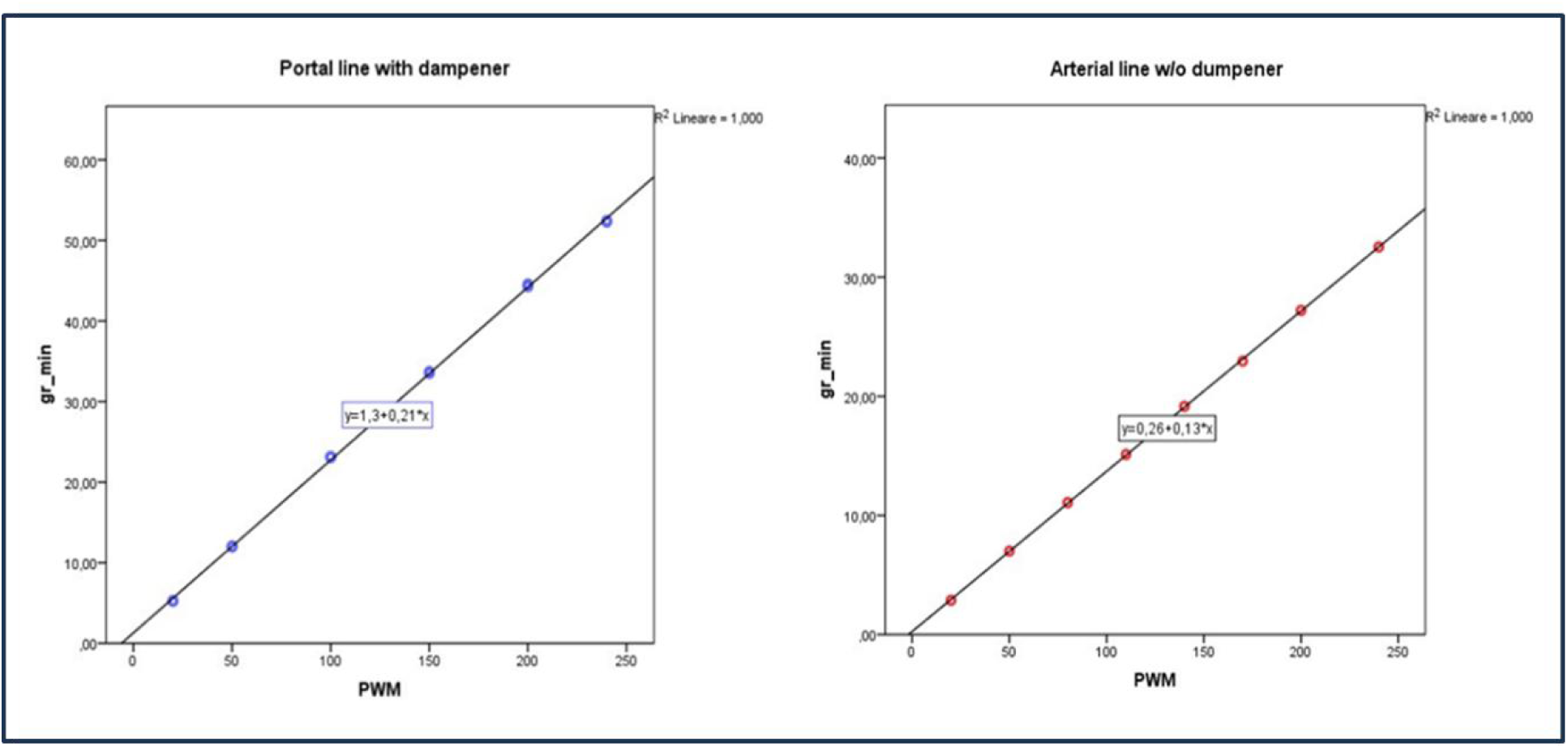
Correlation between the PWM signals and the flow generated by the peristaltic pumps with 3 mm inner diameter tube with the interposition of the dampener2 “portal line” (on the left) and flow generated with a 1 mm inner diameter tube “arterial line” (on the right).

### The oxygenator

At 1 liter/ min of 100% oxygen at 500 mmHg, the mini oxygenator is able to generate a 450 mmHg partial oxygen concentration in a normal saline solution (0.9% NaCl) with a flow rate of 30 ml/min.

### The perfusion Chamber

The perfusion system has been arranged and tested for normothermic arterial and portal perfusion of a rat liver using a PTFE catheter of 2 mm inner diameter and 2 cm long introduced in the portal vein.

The flow resistance of the circuit components between the pressure transducer and the liver has been measured as the slope of the pressure/flow diagram obtained at different flow rates at 37°C using normal saline fluid (Figure 16).

**Figure 16.**
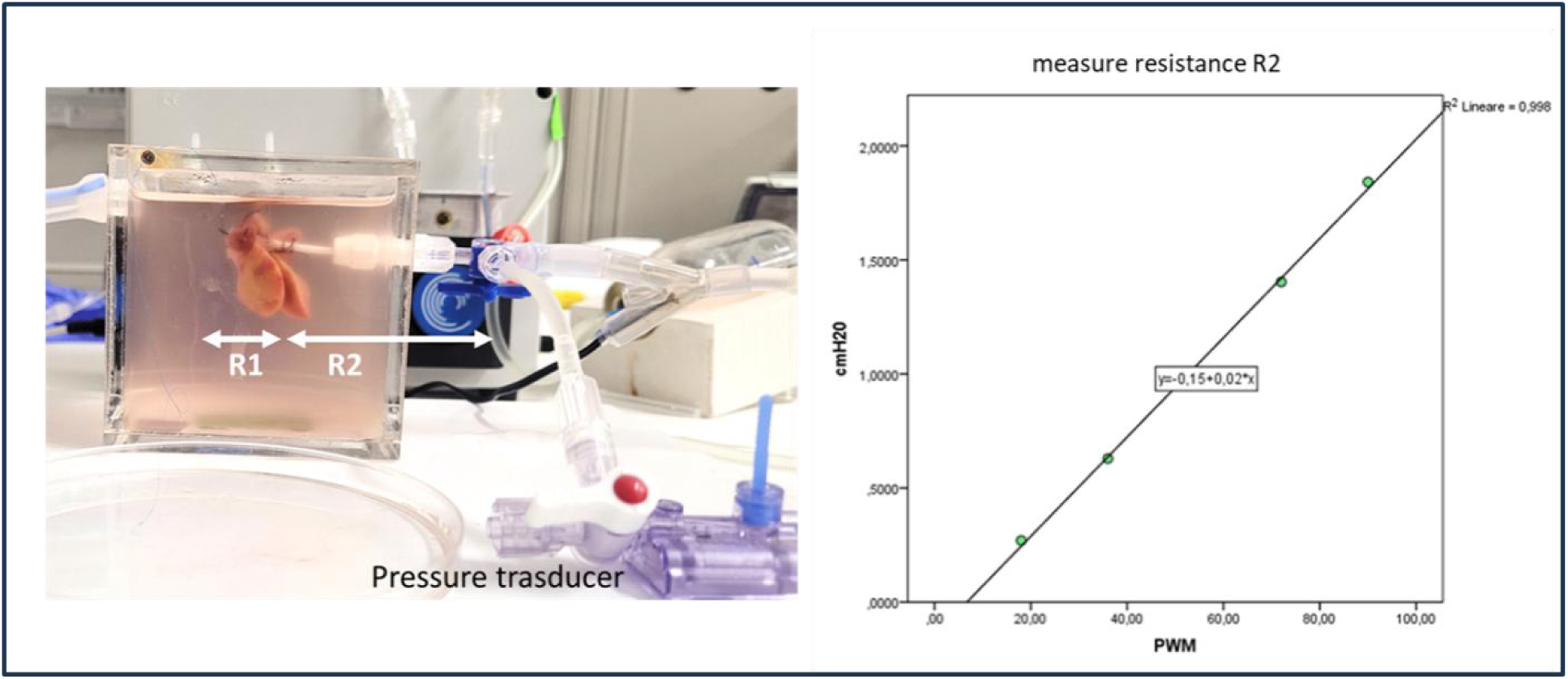
**Left**: A reduced liver rat is displayed during normothermic perfusion. The liver’s resistance (R1) needs to be calculated taking into account the resistance (R2) of the cannula and the connector. **Right**: The R2 value has been calculated as the slope of the pressure/flow line obtained without the liver at different flow-rate.

### Normothermic liver perfusion set-up for compliance study

A simple normothermic set-up (Figure 17) is used in our laboratory to study the compliance variations during the post resection regeneration process in rat livers (data not shown).

**Figure 17.**
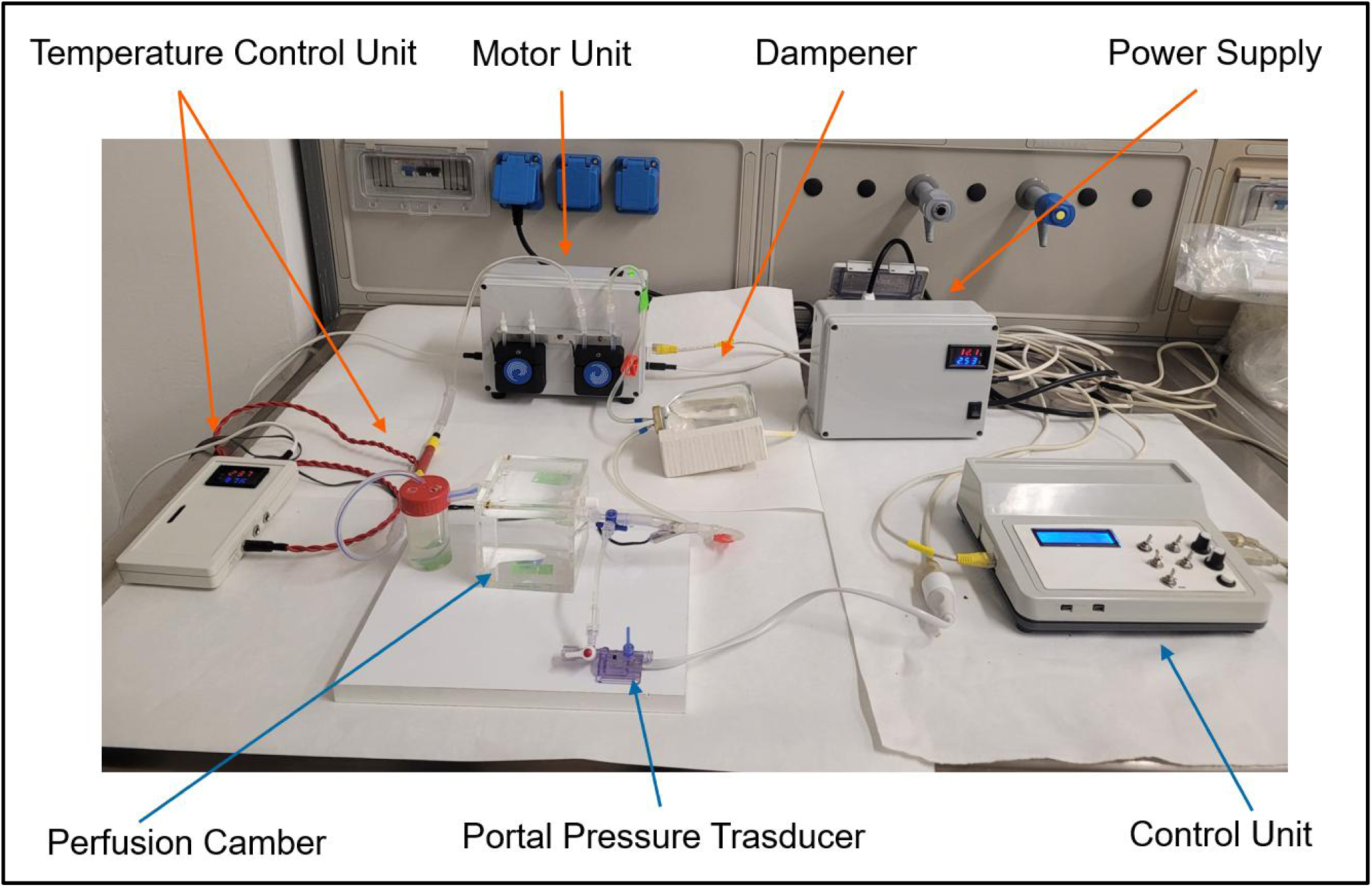
The simplest set-up to obtain flow-pressure data from rat liver by portal vein perfusion.

This simplified circuit allowed to perfuse rat liver through the portal vein at 37 °C at varying flow rates and to measure the portal pression in order to obtain pressure-flow data.

### Hyperthemic perfusion set-up for HIPEC and ILP studies

The modular design enables adaptation to various experimental setups, including peritoneal and limb perfusion (Figure 18). It utilizes peristaltic pumps to regulate inflow and outflow, while pressure transducers and thermocouples provide precise monitoring of the process.

**Figure 18.**
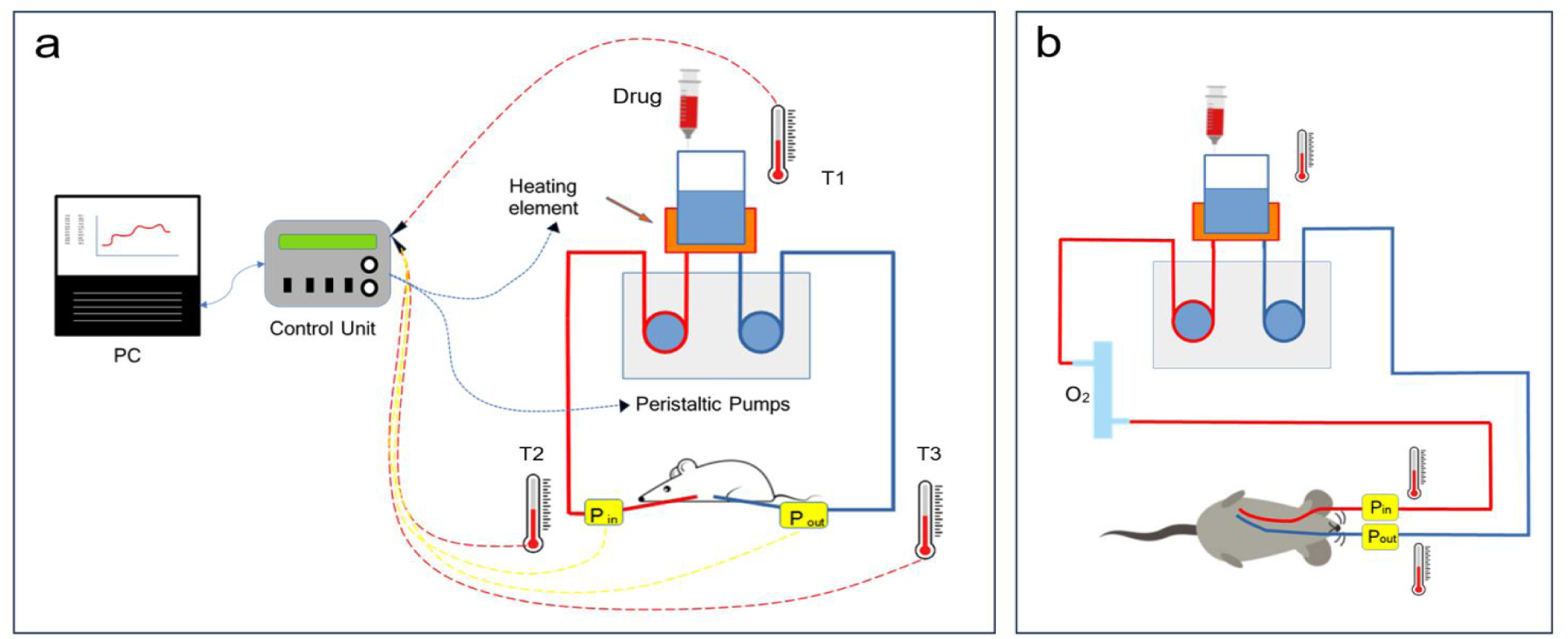
Schematics for Hipec module (a) and for isolated limb perfusion models (b).

A set up for HIPEC models is showed in Figure 19.

**Figure 19.**
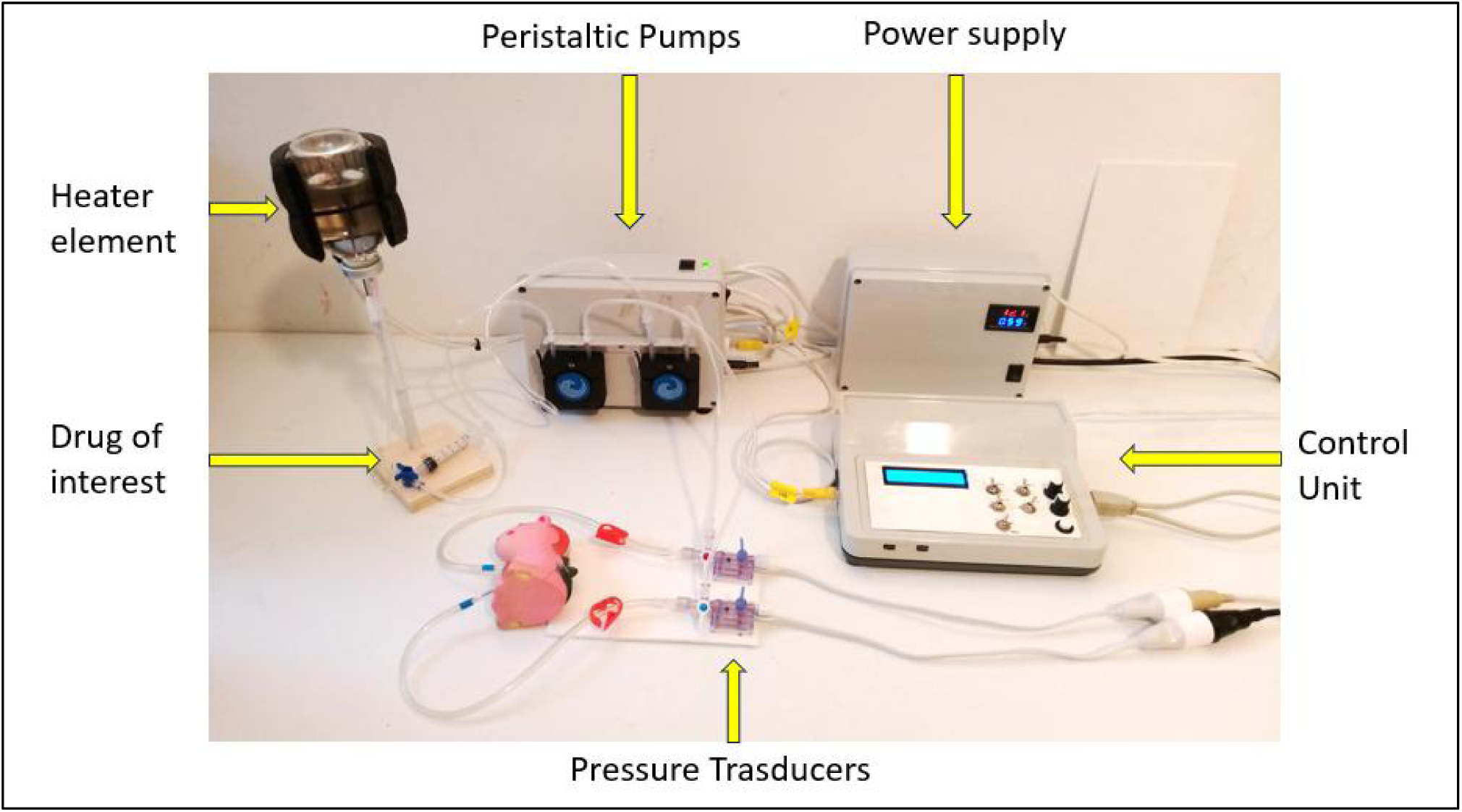
Set up for Hipec module.

### Thermostatic module

The thermostatic unit is designed to heat or cool the organ perfusion liquid in a controlled manner. The fluid from the thermostatic unit circulates in a glass heat exchanger placed in the organ perfusion circuit (Figure 20).

**Figure 20.**
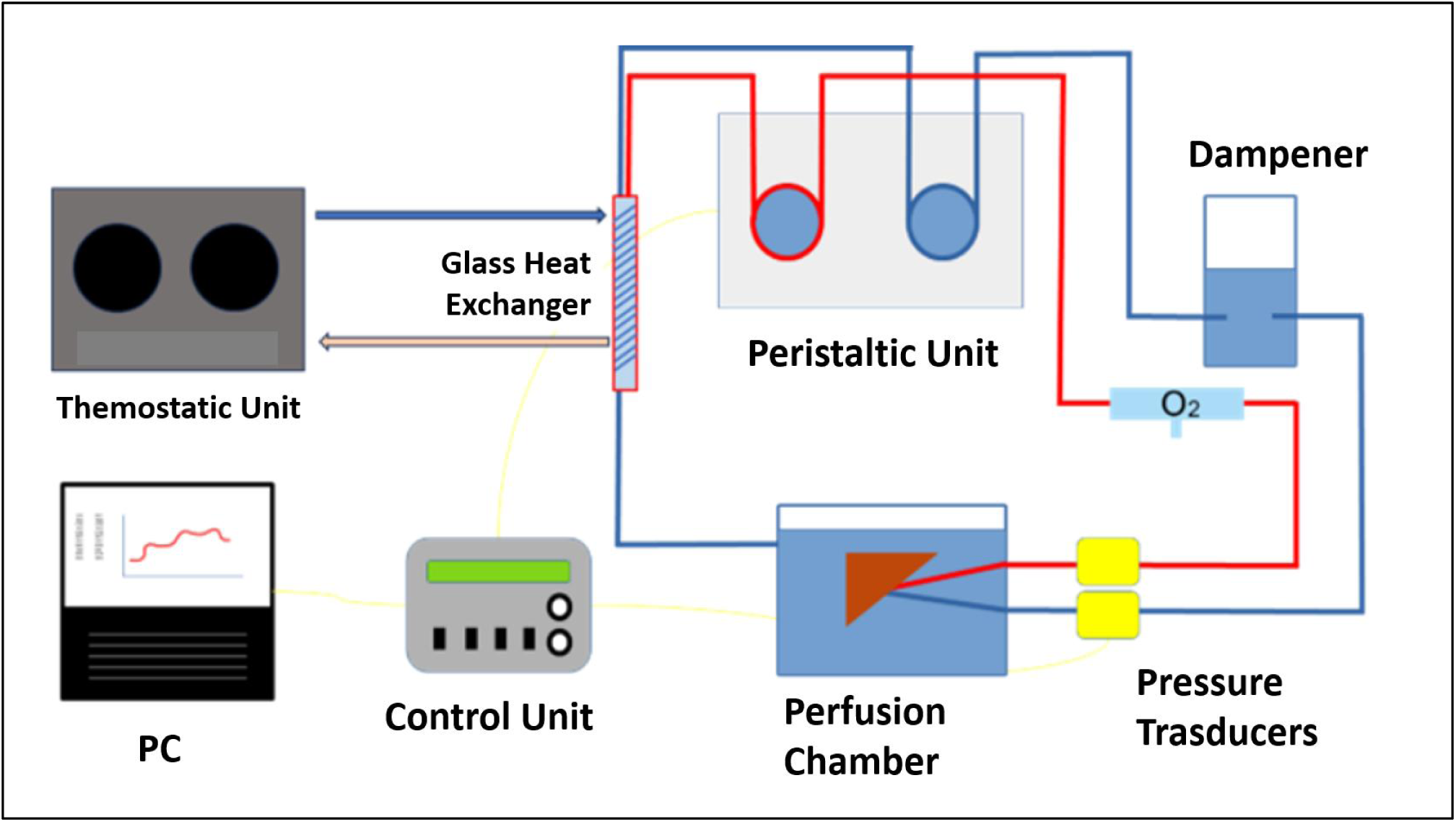
The thermostatic unit is aimed to control the temperature of the perfusion fluid by the interpositio of a glass heat exchanger.

The core component of the device is the Peltier cell, which can heat one side while cooling the other, depending on the polarity of their power supply.

Two 12V Peltier cells are used, each drawing up to 6A: its power supply is regulated through an H-bridge, using a PWM signal generated by an Arduino board (Figure 21).

**Figure 21.**
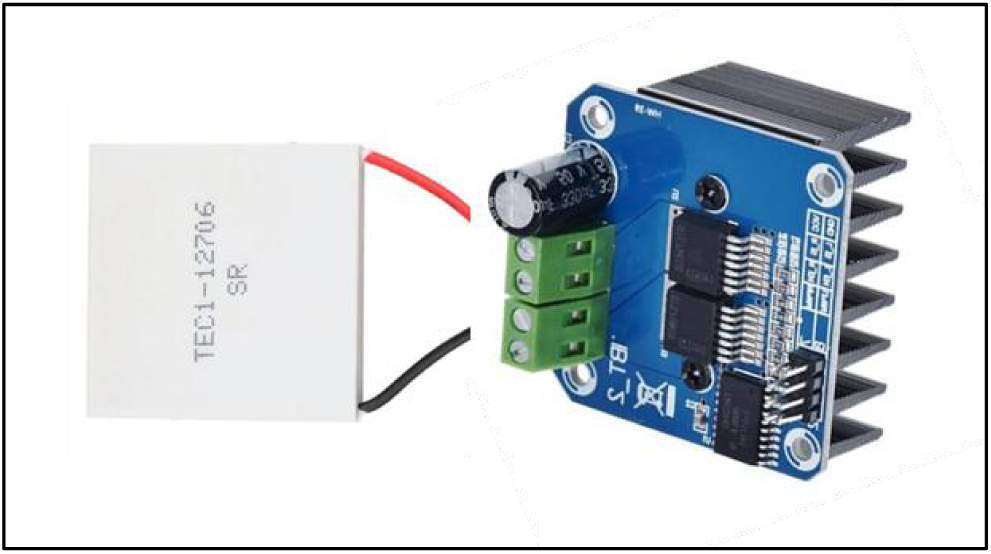
A Peltier cell connected to an H bridge.

**Figure 22.**
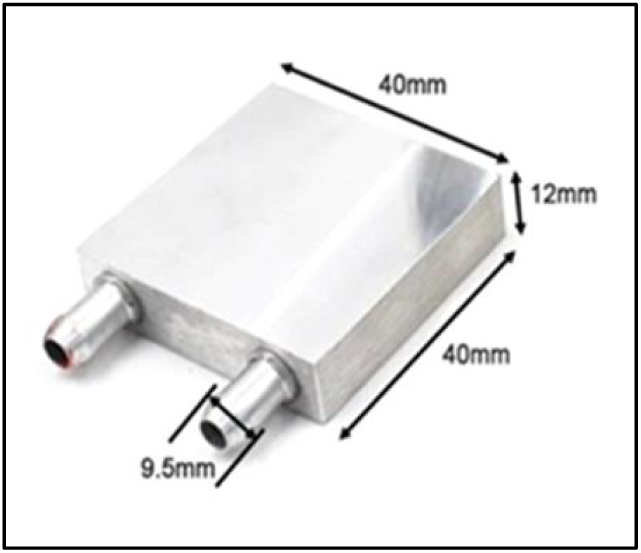
A typical aluminum heat sink.

To work properly, the surfaces of the cell must exchange heat effectively and aluminum heat sinks are used in contact with the Peltier cells (Figure 22).

**Figure 22.**
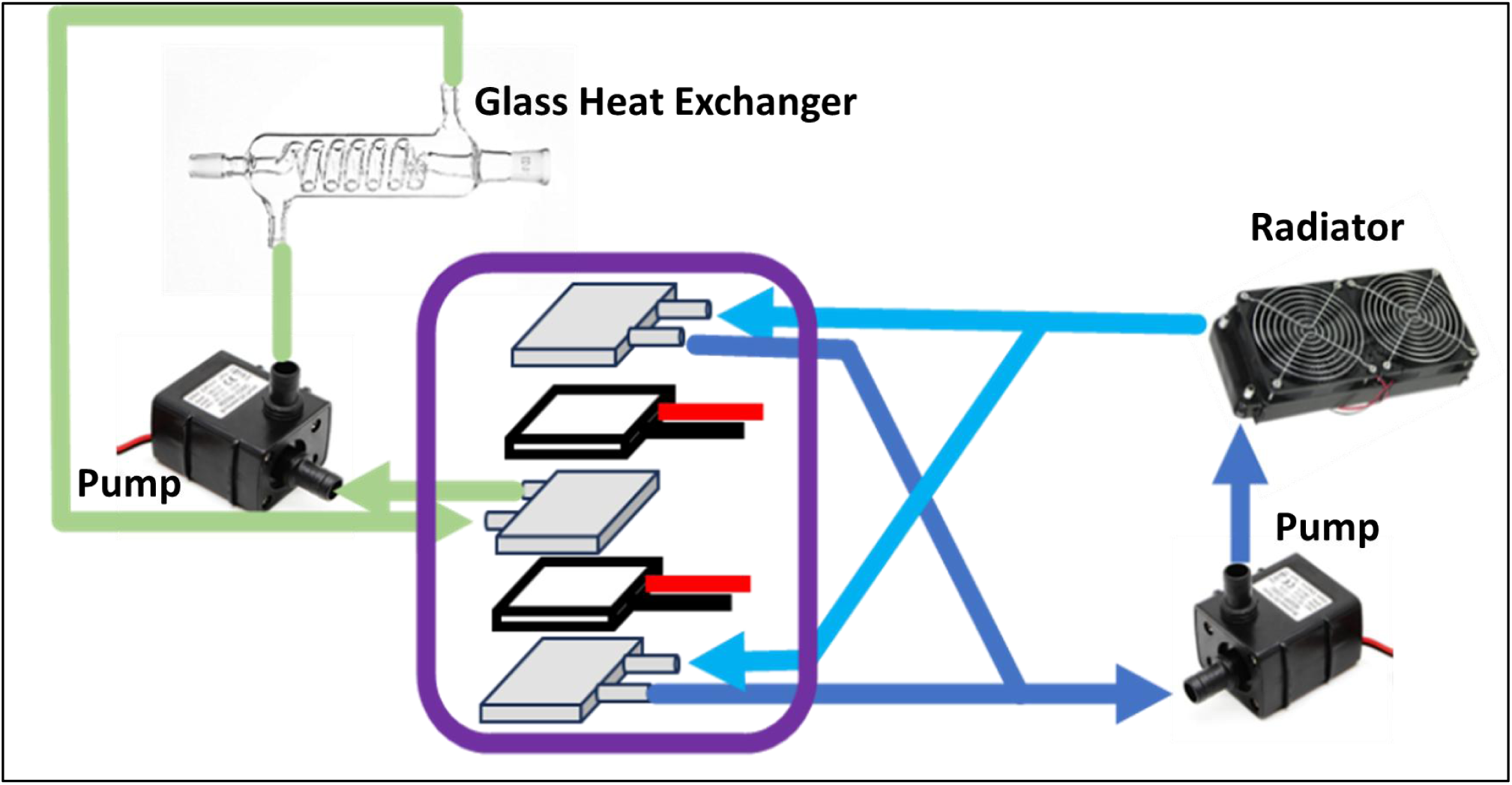
Two Peltier cells are enclosed between aluminum heat sinks (purple circle) creating two indipendet fluid circuits : the green one to the glass heat exchanger and the blue one to the radiator.

There are two independent fluid circuits: one circuit runs through a central heat exchanger situated between the Peltier cells and the glass heat exchanger, while a second one travels through two lateral heat exchangers and then to the radiator; each circuit is powered by its own pump, which is driven by a 12V DC motor (Figure 22).

Negative Temperature Coefficient (NTC) Thermistor are used to monitor the temperature at different site: near the peltier plates, inside the fluid to the glass heat exchanger and inside the perfusion chamber. A voltage divider was build using a 100K NTC : a cubic equation is used for wide range of temperatures, while a linear approximation fits small range (Figure 23).

**Figure 23.**
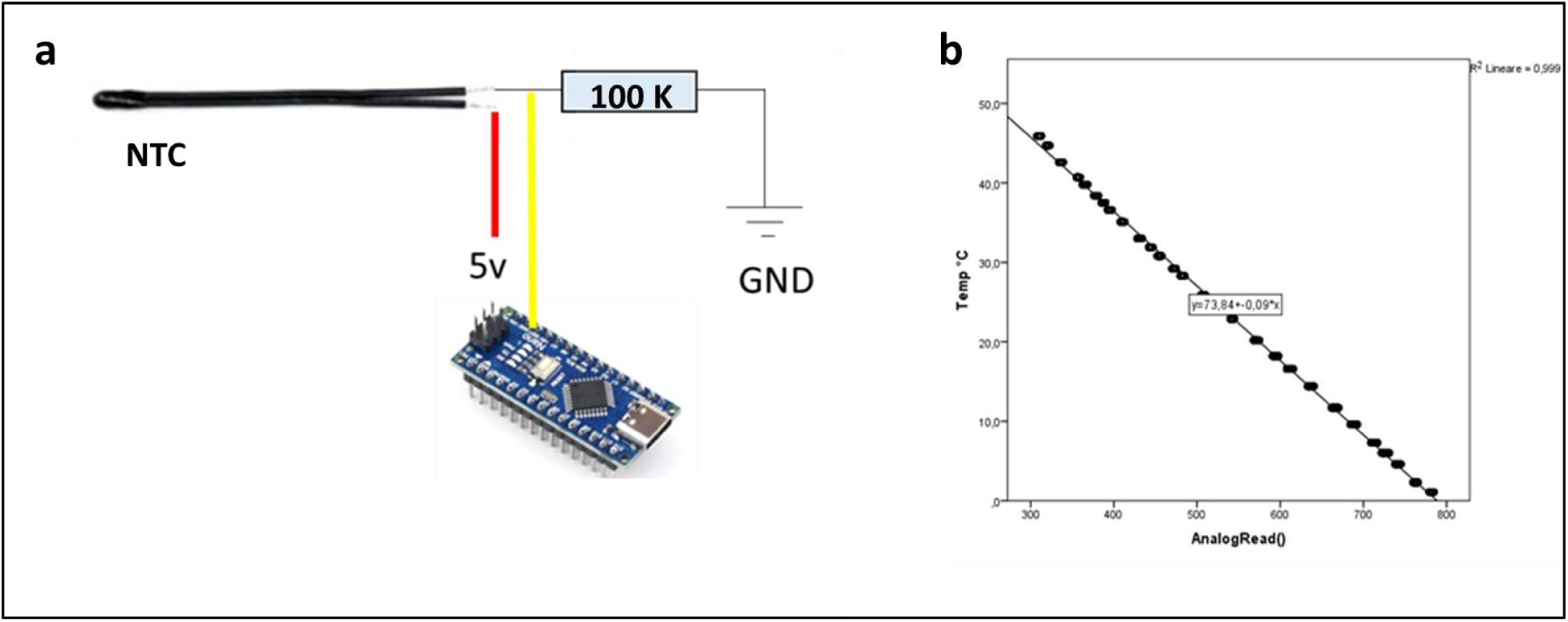
Voltage dividers with the NTC and a 100 kilo-ohm resistor are used to monitor the temperature (a); a linear approximation is used for 0-45 °C measurments (b). The final thermostatic module is showed in figure24.

**Figure 24.**
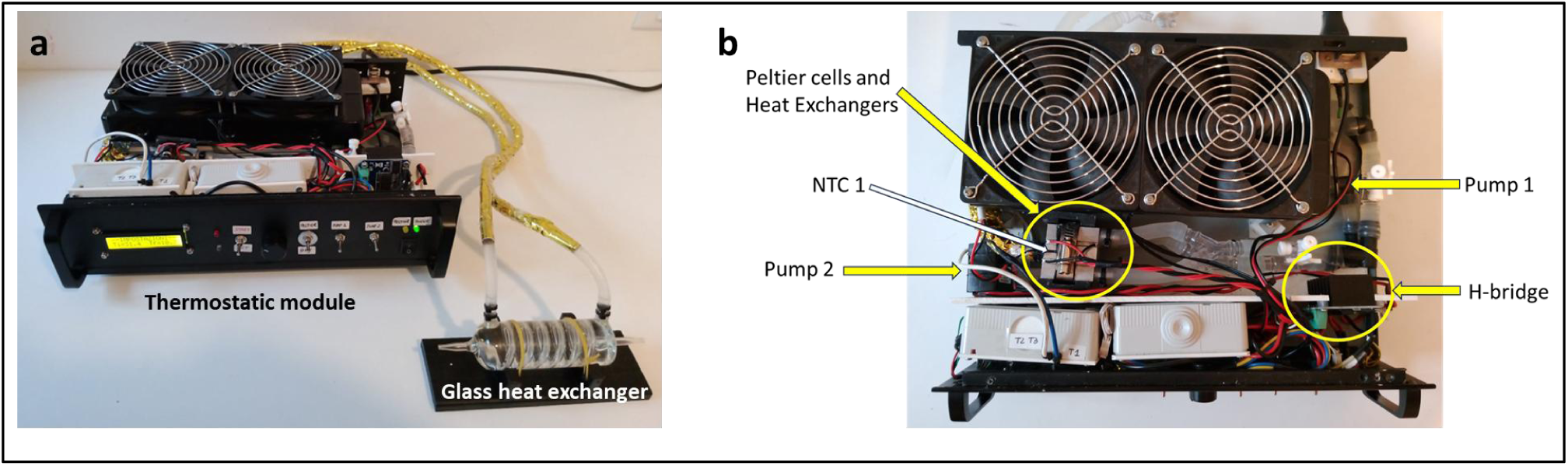
The thermostatic module and the glass heater exchanger are showed in a). The inner configuration with water pumps, peltier cells, aliminum heat sink, one of the NTC probe and the H-bridge is showed in b).

## DISCUSSION

It is common in laboratory settings to assemble experimental procedures using existing devices, often requiring only basic identification, collection, and arrangement of instruments. Occasionally, more invasive modifications of equipment are necessary to meet specific study requirements. However, hacking laboratory instruments is not always straightforward, feasible, or cost-effective, and purchasing new equipment may exceed budgetary constraints.

An alternative approach is the do-it-yourself (DIY) strategy. While designing and building laboratory instruments may seem daunting, particularly for individuals without a background in electronics or computer engineering, platforms like Arduino have democratized this process.

In 2005, the Arduino platform was created at the Interaction Design Institute in Ivrea, Italy, to facilitate the development of interactive prototypes by students lacking expertise in electronics and programming. Arduino is an open-source electronics prototyping platform that interacts with the physical world through sensors, lights, and motors. Over the past decade, it has gained popularity for several reasons: the hardware is open-source and inexpensive; the programming software is user-friendly and free; a wide variety of sensors and shields enhance its flexibility; and a global community of users shares knowledge and projects [10–13].

Like artists, makers, and students, many scientists have embraced the potential of the free and open-source hardware (FOSH) movement, leading to the development of laboratory tools using Arduino [14, 15].

In my experience the process of acquiring the basic skills necessary to program the board and the basic electronic competences was a gradual and pleasant learning experience.

Adapting the pressure sensor transducer and designing the amplifier were the most complex steps, which could be avoided by using Arduino-compatible pressure transducers and amplifier boards readily available on the market.

The final device met the primary objectives of the project: it is user-friendly, cost-effective, and easily modifiable in both hardware and software. The pressure measurements, the flow rate control and the temperature control were sufficiently accurate for our research needs. While formal calibration and accuracy testing were not performed, the device is not intended to replace industrial-grade equipment, which offers superior precision. Nonetheless, the device demonstrated adequate reliability for testing hypotheses in our studies on liver resection in rats.

The system is currently being used in the context of a multidisciplinary basic research project and it is continuously being implemented [16].

In conclusion, the Arduino platform proved to be a pivotal component in developing our perfusion system : its accessibility and versatility made electronics-based projects feasible for our team. Moreover, Arduino and the broader FOSH movements may represent a sustainable and valuable way to enhance scientific education and to increase the opportunities for scientist with limited financial resources.

## Notes

### Competing Interest Statement

The authors have declared no competing interest.

